# Risk Factors Associated to Types of Gallstone Diagnosed at Ibn-Sina Specialized Teaching Hospital, Khartoum, Sudan

**DOI:** 10.1101/640102

**Authors:** Mahasin Ibrahim Shaddad, Azza Abdulrahman Fadl, Ayat Jervase, Nur Ibrahim Ali Garelnabi, Suzan Al Hakem, Tariq Mohamed Hussein, Mohamed H Ahmed, Ahmed Omer Almobark

**Affiliations:** Graduated College, Public and Tropical Health program, University of Medical & Sciences and Technology, Khartoum, Sudan; National Center of Gastrointestinal & Liver Diseases, Ibn-Sina Specialized Teaching Hospital, Khartoum, Sudan; Department of Medicine and HIV Metabolic Clinic, Milton Keynes University Hospital NHS Foundation Trust, Eaglestone, Milton Keynes. Buckinghamshire, UK; Department of Pathology, Faculty of Medicine, University of Medical Sciences and Technology, Khartoum, Sudan

**Keywords:** Cholelithiasis, Gallstone disease, Risk factors, Types of gallstone, Ultrasonography

## Abstract

**Introduction:** Gallstone disease (Cholelithiasis) affects 10 to 15% of the population of developed countries. Our study aimed to determine the risk factors associated to different types of gallstone in patients diagnosed in Khartoum State Ibnsina Gastroenterology Center.

**Methods:** A facility-based prospective cross-sectional study was implemented on a convenient sample of 47 participants diagnosed with gallstone through ultrasonography in Ibnsina Gastroenterology center and who underwent surgical interventions for gallstone removal. A standardized interviewer-administrated research tool comprising three parts was used to collect data related to the characteristics of the participants, their medical presentation and examination as well as information on types of gallstone, surgical interventions and outcomes. The data were computerized through Epi-info^7^ and analyzed through SPSS 23. Descriptive statistics were firstly performed and association was tested through Chi square tests and ANOVA. A multinomial regression analysis established the relationship between types of gallstone and their associated risk factors. All statistical tests were considered as significant when *p* < 0.05.

**Results:** The risk factors statistically associated to gallstone types were family history (*p* = 0.011) and duration of living in the residence area (*p*= 0.043) in pigment-cholesterol gallstone model vs mixed-cholesterol gallstone model. Other four risk factors contributing to the pathogenesis of gallstone were parity (OR = 1.623 [95% CI: 0.795-3.315]) vs (OR = 1.426, [95% CI: 0.729- 2.790]), waist circumference (OR= 1.014 [95% CI: 0.948-1.085]) vs (OR = 1.001 [95% CI: 0.942- 1.065]), chronic disease (OR = 0.698, [95% CI: 0.028 - 17.445]) vs (OR = 0.354, [95% CI: 0.021- 6.087]) and serum triglyceride (OR = 0.985, [95% CI: 0.950- 1.022]) vs (OR= 0.980, [95% CI: 0.949- 1.012]).

**Conclusion:** Our finding indicated six risk factors related to types of gallstone. Further multicenter research in Sudan on risk factors is needed to calibrate and validate our model.

## INTRODUCTION

Gallstone disease (GSD) or cholelithiasis affects 10 to 15% of the population of developed countries [1]. Usually GSD, an asymptomatic disease, is diagnosed incidentally through ultrasonography screening. However, it can present as abdominal/back pain, fever, nausea, vomiting, and jaundice [2].

Various classification systems were proposed to classify gallstones, out of which, the two most popular systems were the National Institute of Health and Japanese Society of Gastroenterology classifications [3]. The National Institute of Health classified gallstones based on their chemical composition and differentiated cholesterol and pigment gallstones. The last one was further subdivided in black and brown gallstones. In the other hand, the Japanese Society of Gastroenterology classified gallstones based on morphology. This classification differentiated cholesterol, pigment and rare gallstones. The cholesterol gallstones were further subdivided in pure cholesterol, combination and mixed gallstones; whereas the pigment gallstones were subdivided in two subtypes (black and calcium bilirubinate gallstones).

The prevalence of the most frequent type of gallstone remained a controversy in the literature. With a prevalence ranging from 36.8 to 53.0% some authors [4–6] reported pure cholesterol gallstone as the predominant type; authors [7–12] reporting a prevalence of pigment gallstone of 37.2% −35% sustained the predominance of this type; elsewhere in the literature, mixed gallstone (66.7% - 89.14%) was reported to be predominate [13–15].

The pathogenesis of GSD is multifactorial: age, gender, ethnicity, tobacco smoking, alcohol consumption, physical activity, diabetes mellitus, metabolic syndrome, non-alcoholic fatty liver disease, hepatitis B and C virus infections, and inflammatory bowel disease [16–27].

A study, conducted on a sample of 102 adult patients who underwent through cholecystectomy, evaluated the etiological risk factors associated to types of gallstone which were classified based on physical characteristic and chemical composition [28]. Black pigment gallstone (47%, 48/102) ranked first followed by mixed cholesterol gallstone (37%, 38/102), pure cholesterol (10%) and brown pigment gallstones (6%). Of the etiological factors assessed for the two first types of gallstone a statistically significant association (*p* =0.018) was found between BMI and types of gallstone as well as between types of gallstone and positive history of type 2 diabetes (*p*=0.035). The other risk factors assessed were family history of GSD, history of dyslipedemia, total cholesterol, triglycerides, high density lipoprotein cholesterol, low density lipoprotein cholesterol, fasting blood glucose, parity and use of exogenous oestrogen in female patients, smoking and alcohol consumption in males were not statistically associated (*p* >0.05) to the types of gallstone. Another study [29] on a sample of 100 patients investigated the association between gallstone characteristics and 23 patient parameters. A statistically significant association was found between both mean diastolic (*p*=0.012) and systolic blood pressure and types of gallstone (*p*=0.027).

The association between diet and types of gallstone was assessed through a matched gender and age case-control study [30] on a sample of 234 patients; 135 cases who benefited from cholecystectomy and 99 controls selected from other patients of the same hospital. The most prevalent was pigment gallstone (43.7%, 59/135) followed by cholesterol (29.6%, 40/135) and mixed (26.7%, 36/135) gallstones. The dietary patterns of cases and controls were regrouped in four factors, namely factor 1 (beef, pork, and fried food), factor 2 (white rice, whole grain, vegetable, and legume), factor 3 (tomato, fruit, and mushroom) and factor 4 (egg, poultry, and seafood). The association between each of those factors and two types of gallstone (pigment and cholesterol gallstones) was determined. Factor 1 was statistically associated with cholesterol gallstone (*p*=0.016), whereas its association with pigment gallstone was not statistically significant (*p*=0.900). The remaining three factors were not statistically associated with both cholesterol and pigment gallstones. The characteristics of the controls and cases of GSD (pigment and cholesterol gallstones) revealed a statistically significant association between gallstone types and family history of GSD (p=0.007) as well as drinking alcohol (*p*=0.002). Age, gender, experience of pregnancy, contraceptive use, hormone replacement therapy, body mass index (BMI), medical history, regular exercise, smoking and used of supplement were all not statistically associated with the two types of gallstone (*p* > 0.05).

The purpose of our research was to determine the risk factors associated to different types of gallstone in patients diagnosed in Khartoum State Ibnsina Gastroenterology Center, Sudan.

## MATERIALS AND METHODS

A facility-based prospective cross-sectional study was implemented in Ibnsina Gastroenterology Center during the period from September to December 2018. The Center has three theater rooms and 28 surgical beds. It performs monthly approximately 250 ultrasonography examinations and 50 surgical interventions. A purposive convenient sampling technique was used to select 47 patients aged ≥18 years regardless their gender, ethnicity and religion. Each of the 47 patients had complete ultrasonography examination records diagnosing GSD; they underwent through surgical intervention for removing their gallstones and had post-surgical follow-up medical records.

A standardized pre-tested interviewer-administrated research tool comprising three parts was used. Variables collected in part 1 were age, gender, occupation, tribe, residence, source of drinking water, alcohol consumption, cigarette smoking, medical history, family history of GSD, gynecological and obstetrical history. Part 1 included also measurements of blood pressure, height, weight, waist circumference; laboratory tests and ultrasonography examination results. Physical and morphological characteristics of gallstones were collected under part 2. The last part of the questionnaire collected variables related to duration of hospitalization, types of surgical intervention and their respective outcomes. The data collected were computerized through a template developed in Epi-info™ 7.1.5.2 and analyzed through SPSS version 23. The data were summarized numerically (mean, standard deviation, median) and graphically (frequency tables for estimating proportions and graphics). Association among categorical variables was tested through chi square tests while association between numerical and categorical variables was assessed through ANOVA. A multinomial regression analysis established the relationship between the types of gallstone and six risk factors (family history of GSD, duration of living in the residence area, parity, chronic disease, serum triglyceride and waist circumference). All statistical tests were considered as statistical significant when *p* < 0.05.

## RESULTS

### Characteristics of the Study Participants

#### Sociodemographic Characteristics of the Study Participants

The majority of the study participants (n=47) were females (89.4%, 42/47); males were 10.6% (5/47). Their median age of 45 years ranged from 19 to 80 years. 72.4% (34/47) of the participants were married, the remaining 27.6% were widows (10.6%), single (8.5%) and divorced (8.5%). The participants were predominately housewives (70.3%, 33/47), 80.9% (38/47) lived in urban areas and 19.1% resided in rural areas with an average years of living in their respective residential area of 30 years ranging from < 1 to 80 years (table 1).

**Table 1:** Characteristics of the study participants (n=47)

| Variable | Number | % |
| --- | --- | --- |
| **Gender (n=47)** |  |  |
| Male | 5 | 10.6 |
| Female | 42 | 89.4 |
| **Age in years (n=47)** |  |  |
| Median | 45 |  |
| Min-Max | 19-80 |  |
| **Marital Status (n=47)** |  |  |
| Married | 34 | 72.4 |
| Widow | 5 | 10.6 |
| Single | 4 | 8.5 |
| Divorced | 4 | 8.5 |
| **Occupation (n=47)** |  |  |
| Housewife | 33 | 70.3 |
| University Student | 4 | 8.5 |
| Teacher | 3 | 6.4 |
| Daily Labor | 2 | 4.3 |
| Farmer | 1 | 2.1 |
| Public Health Officer | 1 | 2.1 |
| Secretary | 1 | 2.1 |
| Shop Owner | 1 | 2.1 |
| Unemployed | 1 | 2.1 |
| **Residence area (n=47)** |  |  |
| Urban | 38 | 80.9 |
| Rural | 9 | 19.1 |
| **Duration of living in the residence area (years)** |  |  |
| Median | 30 |  |
| Min-Max | 0*-80 |  |

\*<1 year.

#### Personal Habits of the Study Participants

The *sources of drinking water* available to the study participants were tape and mineral water (regrouped as “safe source” of drinking water), borehole, “hafeer”, and river (classified as “unsafe source”). In the overall, 61.7% (29/47) of the participants had access to safe source of water and 38.3% (18/47) used unsafe source. A statistically significant association (*p*=0.000) was found between the type of residential area and source of drinking water.

Of the 47 participants, 4.3% (2/47) consumed *alcohol* and the remaining 95.7% (45/47) did not. Unfortunately, information on the frequency and the quantity of alcohol consumed was not recorded. Regarding *cigarette smoking*, of the 47 participants, 91.5% (43/47) had never smoked and 8.5% (4/47) reported to be smokers with 6.4% (3/47) who stopped smoking and 2.1% (1/47) still smoking at the time of the data collection. The four smokers were all males living in urban area with an average year of smoking of 21 years ranging from 14 to 30 years and a daily consumption of 2 packs of ten cigarettes ranging between 1 to 2 packs/day

#### Female Reproductive Characteristics

The *age at menarche* reported by the study participants (n=42) ranged from 8 to 15 years with a median age of 12 years. The mean age at menarche was higher (12.3 years ± 2.2) in females from rural areas than in those living in urban areas (11.8 years ± 1.4); however, this difference was not statistically significant (*p=* 0.502).

Regarding the *gynecological and obstetrical history*, the participants reported an average of 5 pregnancies (range:0-13) and 4 deliveries (range:0-11). When asked if they were *menopause*, half (50.0%, 21/42) of the participants were already menopause and 50.0% were not yet. The median year of those menopause was 7 years ranging from < 1 to 22 years. 64.3% (27/42) were not under *contraception* and 35.7% (15/42) used contraceptive at the time of the study. Combined oral contraceptive pill was the most frequent type (73.3%, 11/15) used.

*Polycystic ovary syndrome* (PCOS) was present in 9.5% (4/42) of the study participants and absent in 90.5% (38/42). In those suffering with PCOS, the median duration of the condition was 8 years varying from 1 to 20 years. A statistically significant association was found between number of pregnancy (*p*=0.023), number of parity (*p=*0.009) and presence of PCOS.

### Medical Presentation of the Participants

#### Symptoms Reported by the Participants

At the time of the data collection, of the 47 participants, *abdominal pain* was reported by 93.6% (44/47). This pain was lasting for an average of 9 months ranging from 1 to 72 months. The predominant pain (63.6%, 28/44) was “right hypochondrial pain associated to epigastric pain” followed by “right hypochondrial pain” and “epigastric pain” with respectively 20.5% (9/44) and 15.9% (7/44). 72.7% (32/44) of the participants related their pain to meals, with 43.8% (14/32) experiencing pain after meals. Shoulder pain prevailed in 44.7% (21/47) of the participants, the same proportion had a history of biliary colic (44.7%, 21/47), fever and jaundice were reported respectively by 25.5% (12/47) and 36.2% (17/47). 10 participants (90.9%, 10/11) reported abdominal distension and one complained of abdominal distension.

#### Chronic Diseases Reported by the Study Participants

Hypertension was the most frequent (19.1%, 9/47) disease harbored by the study participants and the condition lasted from < 1 year to 12 years with a median duration of 3 years. Diabetes mellitus ranked second with a prevalence of 10.6% (5/47) and diabetic participants were living with their condition for an average of 5 years (range: 1-20 years). Hypothyroidism and renal stone ranked in third position with a prevalence of 4.3% (2/47) and an average duration of respectively 4 years and 5.5 years ± 4.9. Ulcerative colitis (2.1%) and cerebrovascular accident (2.1%) were reported respectively by one participant.

#### Family History of Gallstone Disease

The participants were asked if a member of their respective family experienced a gallstone disease, 40.4% (19/47) reported a family history and 59.6% (28/47) did not have a family history of GSD. 19.1% (9/47) of the participants had a first degree family member with a history of GSD, 17.0% (8/47) had a second degree family member affected by GSD and two participants (4.3%, 2/47) had both first and second degree family members who experienced a GSD.

### Medical Examination

#### Physical Examination Results of the Study Participants

This examination comprised the blood pressure and anthropometric measurements of the participants (n=47). The systolic blood pressure of the participants ranged from 100 to 167 mmHg with a median of 126 mmHg and their median diastolic blood pressure was 80 mmHg (range: 60-100 mmHg). Their mean height of 1.6m ±0.1 ranged from 1.3 to 1.8 m and their median body weight of 70 kg ranged from 46 to 117 kg; hence, a median body mass index (BMI) of 26.8 varying from 17.6 to 45.7. The 47 participants had a median waist circumference of 94 cm (range: 39-140 cm).

#### Abdominal Ultrasonography Findings

These findings were summarized in table 2 in three groups: gallbladder, common bile conduct and other ultrasonography findings.

**Table 2:** Results of the ultrasonography examination of the participants (n=47)

| Variable | Number | % |
| --- | --- | --- |
| **1. Gallbladder** |  |  |
| **Gallbladder Wall (n=47)** |  |  |
| Thick | 26 | 55.3 |
| Normal | 21 | 44.7 |
| **Gallbladder Volume (n=47)** |  |  |
| Contracted | 17 | 36.2 |
| Normal | 15 | 31.9 |
| Distended | 15 | 31.9 |
| **Gallstone Number (n=47)** |  |  |
| Multiple | 39 | 83.0 |
| Single | 6 | 12.8 |
| Double | 2 | 4.2 |
| **Gallstone Size (n=47)** |  |  |
| Small | 18 | 38.2 |
| Medium | 17 | 36.2 |
| Large | 6 | 12.8 |
| Sludge | 6 | 12.8 |
| **2. Common Bile Duct** |  |  |
| **CBD Stones (n=47)** |  |  |
| Absent | 34 | 72.3 |
| Present | 13 | 27.7 |
| **CBD Diameter (n=47)** |  |  |
| Normal | 34 | 72.3 |
| Dilated | 13 | 27.7 |
| **Dilated CBD Diameter in mm (n=13)** |  |  |
| Mean (SD) | 12.9 (3.4) |  |
| Median | 13 |  |
| Min-Max | 8-20 |  |
| **3. Other ultrasonography findings** |  |  |
| **Fatty Liver (n=47)** |  |  |
| Absent | 42 | 89.4 |
| Present | 5 | 10.6 |
| **Renal Stones (n=47)** |  |  |
| Absent | 46 | 97.9 |
| Present | 1 | 2.1 |

Regarding the *gallbladder*, the ultrasonography revealed that 55.3% (26/47) of the participants had thick gallbladder wall and the wall was normal for the remaining 44.7% (21/47). The gallbladder volume was contracted in 36.2% (17/47) of the participants and was normal or distended in respectively 31.9% (15/47). Concerning the number of stones diagnosed, the most frequent (83.0%, 39/47) was multiple stones followed by single (12.8%, 6/47) and double stones (4.2%, 2/47). The size of stones varied from small (38.2%, 18/47) to large/sludge (12.8%, 6/47) as revealed by table 2.

In the *common bile duct* (CBD), stones were present in 27.7% (13/47) of the study participants and absents in the remaining 72.3% (34/47). The diameter of CBD was normal in 72.3% of the participants and dilated in 27.7%. For those with dilated CBD, the median diameter of 13 mm ranged from 8 to 20 mm.

The *other ultrasonography findings* were fatty liver and renal stones present respectively in 10.6% (5/47) and 2.1% (1/47) of the participants; the renal stone was diagnosed in the right kidney.

#### Blood Examination Results of the Study Participants

For each of the study participants, the blood parameters investigated were complete blood count (CBC), diabetes mellitus and lipid profile. Regarding the CBC, the white cells counts varied 3.8 to 20.0 ×103/ ul with a mean of 7.1 x 103/ ul ± 3.2; the average red cells counts of 4.4 milli/ul ± 0.6 ranged from 2.7 to 5.7 × 103/ ul. The participants had a mean hemoglobin of 12.5 g/dl ± 1.4 and a mean corpuscular hemoglobin of 28.7 Pg; the mean corpuscular volume was 84.1 fl ± 5.2. Diabetes mellitus was evaluated based on measurements of fasting blood glucose and hemoglobinA1c levels. Their means were respectively 110.9 mg/dl ± 33.6 and 5.3% ± 1.3. The lipid profile of the participants reveled a mean serum cholesterol of 177.9 mg/dl±34.4, a mean serum triglyceride of 98.9 mg/dl±41.8, a mean high density lipoprotein of 50.2 mg/dl± 12.2 and a mean low density lipoprotein of 97.0 mg/dl ±28.3.

#### Surgical Interventions and Outcomes

The most frequent type of surgical intervention performed was the mini-cholecystectomy (53.2%, 25/47), followed by open common bile duct exploration combined with cholecystectomy (27.7%, 13/47) and laparoscopic cholecystectomy (19.1%, 9/47). The hospitalization of the patients lasted on the average (median) 2 days ranging from 2 to 13 days. 93.6% (44/47) fully recovered and 3 patients (6.4%) presented complications (delayed recovery from anesthesia, cardiac arrhythmia after surgery and surgical wound infection).

### Different Types of Gallstone

The predominant (44.7%, 21/47) type of gallstone was mixed gallstone followed by pigment gallstone (40.4%, 19/47) and cholesterol gallstone (14.9%, 7/47). The number of stones found by type of gallstone were recorded as “single”, “double” and “multiple”. All the three quantities were found across the identified types of gallstone (table 3), with multiple stones prevailing more in mixed gallstone.

**Table 3:** Gallstone types and their physical characteristics (n=47)

| Physical Characteristics | Gallstone Type Morphological Classification: number (%) |  |  |  |
| --- | --- | --- | --- | --- |
|  | Cholesterol Stone | Pigment Stone | Mixed Stone | Total |
| **Stone Shape (n=47)** |  |  |  |  |
| Rounded | 4 (57.1%) | 8 (42.1%) | 16 (76.2%) | 28 |
| Faceted | 3 (42.9%) | 4 (21.1%) | 3 (14.3%) | 10 |
| Irregular | 0 (0.0%) | 7 (36.8%) | 2 (9.5%) | 9 |
| **Stone Color (n=47)** |  |  |  |  |
| Black/Blackish brown | 0 (0.0%) | 19 (100.0) | 0 (0.0%) | 19 |
| Greenish/Red/Brownish yellow | 0 (0.0%) | 0 (0.0%) | 21 (100.0%) | 21 |
| White/Pale yellow | 7 (100.0) | 0 (0.0%) | 0 (0.0%) | 7 |
| **Stone Surface (n=47)** |  |  |  |  |
| Smooth | 5 (71.4%) | 8 (42.1%) | 10 (47.6%) | 23 |
| Rough | 2 (28.6%) | 11 (57.9%) | 11 (52.4%) | 24 |
| **Stone Character (n=47)** |  |  |  |  |
| Hard | 6 (85.7%) | 12 (63.2%) | 18 (85.7%) | 36 |
| Soft | 1 (14.3%) | 7 (36.8%) | 3 (14.3%) | 11 |
| **Stone Number (n=47)** |  |  |  |  |
| Single | 1 (14.3%) | 4 (21.0%) | 3 (14.3%) | 8 |
| Double | 1 (14.3%) | 1 (5.3%) | 2 (9.5%) | 4 |
| Multiple | 5 (71.4%) | 14 (73.7%) | 16 (76.2%) | 35 |
| **Median Stone weight in g (Min-Max)** | 0.09 (0.05 - 3.70) | 0.30 (0.02 - 2.90) | 0.60 (0.10 - 4.60) |  |
| **Median stone size in cm** | 0.5 x 0.5 | 0.9 x 0.6 | 1.0 x 0.9 |  |

The weight and the size of stone were recorded respectively in gramme and centimeter. The median weight of the stones was higher (0.60 g; range: 0.10-4.60g) in mixed gallstone; it was 0.30 g (0.02-2.90 g) in pigment gallstone and the lowest (0.09 g; range: 0.05-3.70 g) was in cholesterol gallstone. The median size of stone by type correlated with the respective stone weight with respectively 1.0 cm x 0.9 cm (mixed stone), 0.9 cm x 0.6 cm (pigment stone) and 0.5 cm x 0.5 cm (cholesterol stone). Table 3 displayed the other physical characteristics of the stones by type.

### Types of Gallstone and their Associated Risk Factors

A multinominal regression analysis was performed to assess the relationship between types of gallstones and their associated risk factors. The explanatory variables were family history of GSD, parity, duration of living in the residence area, suffering from a chronic disease, level of serum triglyceride and waist circumference. The regression model fit perfectly the equation with a *p-value* of 0.012.

The types of gallstone were divided in three categories: cholesterol gallstone (n=6, 14.3%), pigment (n=16, 38.1%) and mixed gallstones (n=20, 47.6%). The cholesterol gallstone was used as reference population in the equation.

#### Pigment gallstone relative to cholesterol gallstone

A highly statistically significant association was found between family history of GSD and pigment gallstone with a *p-value* of 0.011, duration of living in the residence area was also statistically (*p*=0.043) associated with pigment gallstone. Despite not statistically significant (*p*=0.184) parity contributed to the model by 1.6 times [95% CI:0.795-3.315] as well as waist circumference with an OR =1.014 [95% CI:0.948-1.085, *p*=0.683]. Serum triglyceride (OR = 0.985, [95% CI:0.950-1.022]; *p*=0.427) and chronic disease (OR = 0.698, [95% CI: 0.028 - 17.445]; *p*=0.826) contributed also to the model at a less extend.

#### Mixed gallstone relative to cholesterol gallstone

None of the six factors of interest was statistically significant (table 4); however, parity contributed to the model by 1.4 times (OR: 1.426, [95% CI: 0.729-2.790]; *p*=0.300). Duration of living in the residence area contribute to the model by 1.1 time (OR = 1.112, [95% CI: 0.980-1.262]; *p*=0.101). Family history of GSD contributed to the model with a coefficient of −3.111(OR = 0.045, [95% CI: 0.002-1.295]; *p*=0.070) representing a contribution of 3 folds. Chronic disease, serum triglyceride and waist circumference contributed to the model at a lesser extent with a coefficient of respectively −1.038 (OR = 0.354, [95% CI: 0.021- 6.087]; *p*=0.474), −0.021 (OR = 0.980, [95% CI: 0.949- 1.012]; *p*=0.212) and 0.001(OR = 1.001, [95% CI: 0.942- 1.065]; *p*=0.963).

**Table 4:** Multinomial regression model estimating the risk factors by types of gallstone with cholesterol gallstone as reference category (n=42).

| Gallstone Type | Variable | B | Std. Error | Wald | df | P | 95% CI for OR |  |  |
| --- | --- | --- | --- | --- | --- | --- | --- | --- | --- |
|  |  |  |  |  |  |  | OR | Lower | Upper |
| **Pigment** | Intercept | -1.096 | 3.524 | 0.097 | 1 | 0.756 |  |  |  |
|  | Family history of GSD | -4.982 | 1.96 | 6.46 | 1 | 0.011 | 0.007 | 0.000 | 0.320 |
|  | Parity | 0.484 | 0.364 | 1.766 | 1 | 0.184 | 1.623 | 0.795 | 3.315 |
|  | Duration of living in the residence area | 0.138 | 0.068 | 4.096 | 1 | 0.043 | 1.147 | 1.004 | 1.311 |
|  | Chronic Disease | -0.360 | 1.643 | 0.048 | 1 | 0.826 | 0.698 | 0.028 | 17.445 |
|  | Serum Triglyceride | -0.015 | 0.019 | 0.63 | 1 | 0.427 | 0.985 | 0.950 | 1.022 |
|  | Waist Circumference | 0.014 | 0.034 | 0.166 | 1 | 0.683 | 1.014 | 0.948 | 1.085 |
| **Mixed** | Intercept | 2.234 | 2.999 | 0.555 | 1 | 0.456 |  |  |  |
|  | Family history of GSD | -3.111 | 1.719 | 3.275 | 1 | 0.070 | 0.045 | 0.002 | 1.295 |
|  | Parity | 0.355 | 0.342 | 1.076 | 1 | 0.300 | 1.426 | 0.729 | 2.790 |
|  | Duration of living in the residence area | 0.106 | 0.065 | 2.689 | 1 | 0.101 | 1.112 | 0.980 | 1.262 |
|  | Chronic Disease | -1.038 | 1.451 | 0.511 | 1 | 0.474 | 0.354 | 0.021 | 6.087 |
|  | Serum Triglyceride | -0.021 | 0.016 | 1.56 | 1 | 0.212 | 0.980 | 0.949 | 1.012 |
|  | Waist Circumference | 0.001 | 0.031 | 0.002 | 1 | 0.963 | 1.001 | 0.942 | 1.065 |

***In the overall***, the risk factor statistically associated to the type of pigment and cholesterol gallstones was family history of GSD contributing by 4 times for pigment and cholesterol gallstones (*p*=0. 011). Duration of living in the residence area was also a statistically significant factor in determining the type of gallstone with a *p-value* of 0.043 for pigment and cholesterol gallstones. The remaining risk factors, not statistically significant, contributed to the pathogenesis of the three types of gallstone with a coefficient ranging from −1.038 (chronic disease) to 0.484 (parity) (table 4).

## DISCUSSION

In Sudan, except a publication [8] reporting age as a risk factor related to different types of gallstone, the available publications reviewed focused either on the chemical constitution [9,10] of gallstone or the surgical technique applied to manage GSD [31,32]. In our study, the participants (n=47) were predominately females (89.4%) with a median age of 45 years (range: 19-80 years). The predominance of female population affected and the age of occurrence of GSD were in the range published by both Sudanese [8–10, 31] and authors from elsewhere [1,33]. However, Adam M.E. et al. agreeing on female paying high tribute to GSD, pointed out the occurrence at an earlier mean age of 31.5 years with 41.7% aged 20-40 years [32].

Premenopausal women are two times at risk to develop GSD compared to males of the same age group [1] this pattern can be attributed to female reproductive hormones and life style [16]. Estrogen can affect gallbladder motility, hence could increase susceptibility to cholesterol gallstone formation [34]. Elder population are 4 to 10 times more likely to developed gallstone with susceptibility to pigment gallstone [33] compared to younger [35]

In our research, the most prevalent type of gallstone was mixed stone (44.7%) followed by pigment stone (40.4%) and cholesterol stone (14.9%). Our findings were in line with published results [13–15] which reported mixed stone as the most frequent. Elsewhere in the literature, pigment stone was the most prevalent type reported [7–12]. Cholesterol stone was published to be the most prevalent in China, Iraqi and New Zealand studies with respectively 36.8%, 49.3% and 53.0% [4–6].

The limitations related to our research on risk factors associated to types of gallstone were the sample size drawn from one hospital and the sampling technique used to select the study participants (all gallstone patients) and the morphological classification used to categorize the types of gallstone. Nonetheless, the multinomial regression analysis performed indicated that ***family history*** (*p* = 0.011) and ***duration of living in the residence area*** (*p*= 0.043) were statistically associated with the type of gallstone (pigment-cholesterol gallstone model vs mixed-cholesterol gallstone model). Despite a no statistically significant association with the types of gallstone, four risk factors which could contribute to the pathogenesis of gallstone were namely (i) the ***parity*** with an OR = 1.623 [95% CI: 0.795-3.315] in pigment-cholesterol gallstone model vs OR = 1.426 [95% CI: 0.729-2.790] in mixed-cholesterol gallstone model, (ii) ***waist circumference*** (OR= 1.014 [95% CI: 0.948-1.085, *p*= 0.683) vs (OR = 1.001 [95% CI: 0.942-1.065]; *p*= 0.963), (iii) ***chronic disease*** (OR = 0.698 [95% CI: 0.028-17.445]; *p*= 0.826) vs (OR = 0.354 [95% CI: 0.021-6.087]; *p*= 0.474) and (iv) ***serum triglyceride***(OR = 0.985 [95% CI: 0.950-1.022]; *p*=0.427) vs (OR = 0.980 [95% CI: 0.949-1.012]; *p*=0.212). Our attempt to assess the risk factors related to types of gallstone through a multinomial regression was already published by Goktas S.B. et al. [36] who assessed the risk factors associated to pigment and cholesterol gallstones in a sample of 164 participants. They found a statistically significant association between positive family history of GSD (*p*=0.011 for pigment gallstone vs *p*=0.317 cholesterol gallstone) with a OR of 1.68 [95% CI:0.61-4.65], other statistically significant association they pointed out were menopause present (*p*=0.010 vs *p*=0.006), presence of anemia (*p*=0.045 vs *p*=0.043) and presence of liver disease (*p*=0.002 vs *p*=0.001), no milk consumption (*p*=0.050 vs *p*=0.001), olive oil consumption (*p*=0.000 vs *p*=0.000), water consumption < 1 litre (p=0.403 vs *p*= 0.213) with OR = 1.83 [95% CI: 0.71-4.72].

The risk factors associated to types of gallstone remained a debate in the literature [6,8,28–30,37]. A new window in research on pathogenesis of GSD was opened by authors [38–42] who pleaded for investigating genetic and environmental patterns.

We assessed six risk factors related to types of gallstone. Our model was based on family history of GSD, duration of living in the residence area, parity, chronic disease, serum triglyceride and waist circumference. Further multicenter research in various hospitals country-wide is necessary to calibrate and validate our model.

## CONCLUSIONS

The surgical outcomes rates were motivating with 93.6% full recovery and 6.4% complications with no case of death revealing the availability of expertise to manage gallstone in our study setting. The challenges remain to identify the risk factors associated to types of gallstone; the six risk factors identified are not exhaustive and plead for further research to find out all the risk factors related to gallstone types through a multidisciplinary collaborative work between health professionals.

## Funding

Health Insurance Corporation Khartoum State (HICKS), Khartoum, Sudan.

## SUPPORTING INFORMATION

**<u>S1</u>:** Assessment of the association between types of gallstone and its associated factors through Chi-square Likelihood Ratio Test (n=47).

**<u>S2</u>:** Assessment of the association between types of gallstone and its associated factors through ANOVA Test (n=47).

## Data Availability

In accordance to data sharing and policy of bioRxiv on the matter, the authors declared that if the submitted manuscript is accepted for publication the data will be deposited in the generalist repository of Dryad.

## CONFLICT OF INTEREST

No conflict of interest.

## ETHICAL CLEARANCE

The research proposal reviewed and adopted by Sumasri Institutional Review Board of the University of Medical Sciences and Technology was approved by the administration of Ibnsina Gastroenterology Center, Khartoum State, Sudan. A well verbal informed consent was obtained from each of the study participants. They were informed on their right to withdraw from the study at any time they might wish and their confidentiality was ensured through the use of anonymous questionnaire and they were secured that the data collected will not be used for any other purpose than the objectives of the study.

## REFERENCES

1. Stinton L.M, Shaffer E.A. Epidemiology of Gallbladder Disease: Cholelithiasis and Cancer. Gut and Liver 2012, Vol. 6, No. 2, pp. 172–187. https://dx.doi.org/10.5009%2Fgnl.2012.6.2.172.

2. Tazuma S., Unno M., Igarashi Y. et al. Evidence-based clinical practice guidelines for cholelithiasis 2016. J Gastroenterol 2017 52: 276–300. https://doi.org/10.1007/s00535-016-1289-7.

3. Kim I.S., Myung S.J., Lee S.S. et al. Classification and Nomenclature of Gallstones Revisited. Yonsei Medical Journal. Vol. 44, No. 4, pp. 561–570, 2003.

4. Qiao T, Ma R-h, Luo X-b, Yang L-q, Luo Z-l, et al. (2013) The Systematic Classification of Gallbladder Stones. PLoS ONE 8(10): e74887. doi:10.1371/journal.pone.0074887

5. Taher M.A. Descriptive study of cholelithiasis with chemical constituents’ analysis of gallstones from patients living in Baghdad, Iraq. International Journal of Medicine and Medical Sciences Vol. 5(1), pp. 19–23, January 2013. DOI: 10.5897/IJMMS12.031.

6. Stringer M.D., et al. Gallstones in New Zealand: composition, risk factors and ethnic differences. ANZ J Surg 83(2013) 575–580.

7. Das B., Malik A.K., Rehman A. et al. Quantitative Analysis of Chemical Composition of Gallstones in North Indian Population (Rohilkhand Region, Uttar Pradesh). NJIRM 2014; Vol. 5(4). July-August, p.4–12.

8. Idris S.A., Shalayel M. HF., Elsiddig K.E. et al. Prevalence of Different Types of Gallstone in Relation to Age in Sudan. Sch. J. App. Med. Sci., 2013; 1(6):664–667.

9. Idris S.A., Elsiddig K.E., Hamza A.A et.al. Extensive Quantitative Analysis of Gallstones. International Journal of Clinical Medicine, 2014, 5, 42–50. http://dx.doi.org/10.4236/ijcm.2014.51009.

10. Idris S.A., Elsiddig K.E., Hafiz M.M. et.al. Minerals’ composition of different types of gallstones in Sudanese population. Open Science Journal of Analytical Chemistry. Vol. 1, No. 1, 2014, pp.1–5.

11. Sharma R., Soy S., Kumar C. et.al. Analysis of gallstone composition and structure in Jharkhand region. Indian J Gastroenterol 2015. DOI 10.1007/s12664-014-0523-6. https://doi.org/10.1007/s12664-014-0523-6.

12. Weerakoon H, Navaratne A, Ranasinghe S, Sivakanesan R, Galketiya K B, Rosairo S (2015) Chemical Characterization of Gallstones: An Approach to Explore the Aetiopathogenesis of Gallstone Disease in Sri Lanka. PLoSONE 10 (4): e0121537. doi:10.1371/journal.pone.0121537

13. Dattal D. S. et.al. Morphological spectrum of gall bladder lesions and their correlation with Cholelithiasis. Int J Res Med Sci 2017 Mar; 5(3); 840–846. www.msgonline.org. DOI: http://dx.doi.org/10.18203/2320-6012.ijrms20170622.

14. Pradhan S.B., Joshi M.R., Vaidya A. et.al. Prevalence of different types of gallstone in the patients with cholelithiasis at Kathmandu Medical College, Nepal. Kathmandu University Medical Journal (2009), Vol. 7, No. 3, Issue 27, 268–271.

15. Gaharwar A., Mishra S.R., Kumar V. A study on Cholecystectomy Specimens. J. Anat. Sciences, 24 (1): June 2016, 7–12.

16. Figueiredo et al. Sex and ethnic/racial-specific risk factors for gallbladder disease. BMC Gastroenterology (2017) 17: 153. https://doi.org/10.1186/s12876-017-0678-6.

17. Shabanzadeh D.M. et.al. Are incident gallstones associated to sex-dependent changes with age? A cohort study. Andrology, 1–8. https://doi.org/10.1111/andr.12391.

18. Aune D. et al. Tobacco smoking and the risk of gallbladder disease. Eur J Epidemiol (2016) 31: 643–653. DOI 10.1007/s10654-016-0124-z. https://doi.org/10.1007/s10654-016-0124-z.

19. Wang J., Duan X., Li B. et al. Alcohol consumption and risk of gallstone disease: a meta-analysis. European Journal of Gastroenterology & Hepatology 2017, 29: e19–e28. http://dx.doi.org/10.1097/MEG.0000000000000803.

20. Zhang Y. et.al. Physical Activity and the Risk of Gallstone Disease a Systematic Review and Meta-analysis. J Clin Gastroenterol 2017; 51: 857–868. https://doi.org/10.1097/MCG.0000000000000571.

21. Aune, D., & Vatten, L.J., Diabetes mellitus and the risk of gallbladder disease: A systematic review and meta-analysis of prospective studies, Journal of Diabetes and Its Complications (2015), http://dx.doi.org/10.1016/j.jdiacomp.2015.11.012.

22. Daniel Mønsted Shabanzadeh, Lars Tue Sørensen & Torben Jørgensen (2016) Determinants for gallstone formation - a new data cohort study and a systematic review with meta-analysis, Scandinavian Journal of Gastroenterology, 51:10, 1239–1248, DOI: 10.1080/00365521.2016.1182583.

23. Mohamed H. Ahmed, Salma Barakat & Ahmed O. Almobarak (2014) The association between renal stone disease and cholesterol gallstones: the easy to believe and not hard to retrieve theory of the metabolic syndrome, Renal Failure, 36:6, 957–962, DOI: 10.3109/0886022X.2014.900424.

24. Jaruvongvanich V., Sanguankeo A., Upala, S. Significant Association Between Gallstone Disease and Nonalcoholic Fatty Liver Disease: A Systematic Review and Meta-Analysis. Dig Dis Sci. DOI 10.1007/s10620-016-4125-2. https://doi.org/10.1007/s10620-016-4125-2.

25. Wijarnpreechaa K. et al. Hepatitis B virus infection and risk of gallstones: a systematic review and meta-analysis. European Journal of Gastroenterology & Hepatology 2016, 28:1437–1442. https://doi.org/10.1097/MEG.0000000000000754.

26. Wijarnpreecha K, Thongprayoon C, Panjawatanan P, Lekuthai N, Ungprasert P. Hepatitis C virus infection and risk of gallstones: a meta-analysis. J Evid Based Med. 2017;10:263–270. https://doi.org/10.1111/jebm.122277.

27. Zhang F.M, Xu C. F., Shan G. D. et al. Is gallstone disease associated with inflammatory bowel diseases? A meta-analysis. Journal of Digestive Diseases 2015;16; 634–641. https://doi.org/10.1111/1751-2980.12286.

28. Weerakoon et al. Can the type of gallstones be predicted with known possible risk factors?: a comparison between mixed cholesterol and black pigment stones. BMC Gastroenterology 2014 14: 88. https://doi.org/10.1186/1471-230X-14-88.

29. Sharma H., Gupta G., Sharma M. K. Correlation of Gallstone Characteristics with the Cl, Volinical Parameters in Cases of Cholelithiasis. International Journal of Anatomy, Radiology and Surgery, 2015 Jul-4(3):1–5.

30. Park Y. et al. Association between diet and gallstones of cholesterol and pigment among patients with cholecystectomy: case-control study in Korea. Journal of Health, Population and Nutrition (2017) 36:39. https://doi.org/10.1186/s41043-017-0116-y.

31. Mohamed M.I., Abdalla A.A, Alshaikh A. A. et al. Laparoscopic Cholecystectomy: A 15-years Experience at a Single Centre, Wad Medani, Sudan. East Cent. Afr. J. Surg. 2014, Vol. 19, No. 2, p:12–16.

32. Adam M.E., Idris S.A. Open minimally invasive cholecystectomy in Khartoum North Teaching Hospital, Sudan. Sch. J. App. Med. Sci., 2014; 2(1A):121–124.

33. Shaffer E.A. Epidemiology and Risk Factors for Gallstone Disease: Has the Paradigm Changed in the 21st Century? Current Gastroenterology Report 2005, 7:132–140. https://doi.org/10.1007/s11894-005-0051-8.

34. Wang H.H., Liu M., Clegg D.J. et.al. New insights into the molecular mechanisms underlying effects of estrogen on cholesterol gallstone formation. Biochim Biophys Acta. 2009 November; 1791(11): 1037–1047. doi:10.1016/j.bbalip.2009.06.006.

35. Acalovschi M. Cholesterol gallstones: from epidemiology to prevention. Postgrad Med J 2001; 77: 221–229. http://dx.doi.org/10.1136/pmj.77.906.221.

36. Goktas S.B., Manukyan M., Selimen D. Evaluation of Factors Affecting the Type of Gallstone. Indian J Surg (February 2016) 78 (1): 20–26. https://doi.org/10.1007/s12262-015-1313-9

37. Parambil SM, Matad S, Soman KC. Epidemiological, demographic and risk factor profile in patients harbouring various types of gallbladder calculi: a cross sectional study from a south Indian tertiary care hospital. IntSurg J 2017; 4:525–8.

38. Xue P, Niu W-Q, Jiang Z-Y, Zheng M-H, Fei J (2012) A Meta-Analysis of Apolipoprotein E Gene e2/e3/e4 Polymorphism for Gallbladder Stone Disease. PLoS ONE 7(9): e45849. doi:10.1371/journal.pone.0045849

39. Gong Y, Zhang L, Bie P, Wang H (2013) Roles of ApoB-100 Gene Polymorphisms and the Risks of Gallstones and Gallbladder Cancer: A Meta-Analysis. PLoS ONE 8(4): e61456. doi:10.1371/journal.pone.0061456

40. Jiang Z-Y, Cai Q, Chen E-Z (2014) Association of Three Common Single Nucleotide Polymorphisms of ATP Binding Cassette G8 Gene with Gallstone Disease: A Meta-Analysis. PLoS ONE 9(1): e87200. doi:10.1371/journal.pone.0087200

41. Katsika D., Grjibovski A., Einarsson C. Genetics and Environmental Influences on Symptomatic Gallstone Disease: a Swedish Study of 43,141 Twins Paris. HEPATOLOGY, Vol. 41, No. 5, 2005, 1138–1143.

42. Martinez-Lopez E. et al. Influence of ApoE and FABP2 polymorphisms and environmental factors in the susceptibility of gallstone disease. ANNALS of Hepatology, July-August, Vol.14, No.4, 2015:515–523.

