## Supplemental Table for "Risk Factors Associated to Types of Gallstone Diagnosed at Ibn-Sina Specialized Teaching Hospital, Khartoum, Sudan"

|  | **Gallstone types** | | | |  |
| --- | --- | --- | --- | --- | --- |
| **Variable** | **Cholesterol** | **Pigment** | **Mixed** | **Total** | ***p-value*** |
| **Gender (n=47)** |  |  |  |  |  |
| Male | 1 (14.3%) | 3 (15.8%) | 1 (4.8%) | 5 (10.6%) |  |
| Female | 6 (85.7%) | 16 (84.2%) | 20 (95.2) | 42 (89.4%) | 0.473 |
| **Marital Status (n=47)** |  |  |  |  |  |
| Married | 5 (71.4%) | 14 (73.7%) | 15 (71.4%) | 34 (72.3%) |  |
| Other Marital Status | 2 (28.6%) | 5 (26.3%) | 6 (28.6%) | 13 (27.7%) | 0.986 |
| **Occupation (n=47)** |  |  |  |  |  |
| Housewife/unemployed | 4 (57.1%) | 15 (78.9%) | 15 (71.4%) | 34 (72.3%) |  |
| Working | 3 (42.9%) | 4 (21.1%) | 6 (28.6%) | 13 (27.7%) | 0.552 |
| **Residence area (n=47)** |  |  |  |  |  |
| Urban | 7 (100.0%) | 12 (63.2%) | 19 (90.5%) | 38 (80.9%) |  |
| Rural | 0 (0.0%) | 7 (36.8%) | 2 (9.5%) | 9 (19.1%) | 0.021* |
| **Source of Drinking Water (n=47)** |  |  |  |  |  |
| Safe Source of Drinking Water | 7 (100.0%) | 9 (47.4%) | 13 (61.9%) | 29 (61.7%) |  |
| Unsafe Source of Drinking Water | 0 (0.0%) | 10 (52.6%) | 8 (38.1%) | 18 (38.3%) | 0.015* |
| **Toxic habits Cigarette/Alcohol (n=47)** |  |  |  |  |  |
| No | 6 (85.7%) | 16 (84.2%) | 20 (95.2%) | 42 (89.4%) |  |
| Yes | 1 (14.3%) | 3 (15.8%) | 1 (4.8%) | 5 (10.6) | 0.473 |
| **Menopause (n=42)** |  |  |  |  |  |
| No | 5 (83.3%) | 6 (37.5%) | 10 (50.0%) | 21 (50.0%) |  |
| Yes | 1 (16.7%) | 10 (62.5%) | 10 (50.0%) | 21 (50.0%) | 0.141 |
| **Use contraceptive (n=42)** |  |  |  |  |  |
| No | 3 (50.0%) | 11 (68.8%) | 13 (65.0%) | 27 (64.3%) |  |
| Yes | 3 (50.0%) | 5 (31.2%) | 7 (35.0%) | 15 (35.7%) | 0.720 |
| Total | 6 | 16 | 20 | 42 |  |
| **Presence of Chronic Disease (n=47)** |  |  |  |  |  |
| Absent | 3 (42.9%) | 9 (47.4%) | 14 (66.7%) | 26 (55.3%) |  |
| Present | 4 (57.1%) | 10 (52.6%) | 7 (33.3%) | 21 (44.7%) | 0.36 |
| **Family history of GSD (n=47)** |  |  |  |  |  |
| No | 1 (14.3%) | 15 (78.9%) | 12 (57.1%) | 28 (59.6%) |  |
| Yes | 6 (85.7%) | 4 (21.1%) | 9 (42.9%) | 19 (40.4%) | 0.009* |

**S1: Assessment of the association between types of gallstone and its associated factors through Chi-square Likelihood Ratio Test (n=47).**

**S2**: **Assessment of the association between types of gallstone and its associated factors through ANOVA Test (n=47).**

|  | **Types of Gallstone** | | | | | | | | | | | |  |  |  |  | |  | |
| --- | --- | --- | --- | --- | --- | --- | --- | --- | --- | --- | --- | --- | --- | --- | --- | --- | --- | --- | --- |
|  | **Cholesterol** | | | | | **Pigment** | | | | **Mixed** | | | **Sum of Squares of groups** | | **Mean Square of groups** | | |  | |
| **Variable** | **Mean** | | **n** | **Std** | **Mean** | | **n** | **Std** | **Mean** | | **n** | **Std** | **Between** | **Within** | **Between** | **Within** | **F** | | ***p-value*** |
| Age in years | | 41.3 | 7 | 16.6 | 54.6 | | 19 | 16.0 | 45.1 | | 21 | 14.8 | 1308.981 | 10679.870 | 654.491 | 242.724 | 2.696 | | 0.079 |
| Duration of Living in Residence (years) | | 23.0 | 7 | 20.2 | 40.4 | | 19 | 23.3 | 27.2 | | 21 | 19.3 | 2399.790 | 19683.870 | 2399.790 | 19683.870 | 2.682 | | 0.080 |
| Age at Menarche | | 12.8 | 6 | 0.8 | 11.9 | | 16 | 1.7 | 11.7 | | 20 | 1.6 | 6.486 | 97.133 | 3.243 | 2.491 | 1.302 | | 0.284 |
| Number of Pregnancies | | 1.8 | 6 | 1.7 | 6.7 | | 16 | 3.1 | 4.9 | | 20 | 3.4 | 105.072 | 372.071 | 52.536 | 9.540 | 5.507 | | 0.008* |
| Parity | | 1.5 | 6 | 1.6 | 5.4 | | 16 | 2.7 | 4.0 | | 20 | 3.1 | 66.583 | 307.250 | 33.292 | 7.878 | 4.226 | | 0.022* |
| Menopause Duration in years | | 2.0 | 1 | . | 12.4 | | 10 | 6.7 | 6.1 | | 10 | 6.6 | 248.510 | 801.300 | 124.255 | 44.517 | 2.791 | | 0.088 |
| Systolic Blood Pressure (mmHg) | | 131.9 | 7 | 16.2 | 124.5 | | 19 | 14.6 | 126.1 | | 21 | 15.0 | 280.426 | 9931.404 | 140.213 | 225.714 | 0.621 | | 0.542 |
| Diastolic Blood Pressure (mmHg) | | 84.7 | 7 | 7.4 | 77.9 | | 19 | 9.8 | 75.6 | | 21 | 8.2 | 438.873 | 3388.361 | 219.437 | 77.008 | 2.850 | | 0.069 |
| Height in meter | | 1.6 | 7 | 0.1 | 1.6 | | 19 | 0.1 | 1.6 | | 21 | 0.1 | 0.002 | 0.331 | 0.001 | 0.008 | 0.154 | | 0.858 |
| Weight in kg | | 76.7 | 7 | 24.1 | 67.3 | | 19 | 16.8 | 70.0 | | 21 | 10.0 | 451.982 | 10534.486 | 225.991 | 239.420 | 0.944 | | 0.397 |
| Waist Circumference in cm | | 92.7 | 7 | 35.5 | 89.7 | | 19 | 19.8 | 85.6 | | 21 | 20.0 | 331.957 | 22600.256 | 165.979 | 513.642 | 0.323 | | 0.726 |
| Body mass index | | 29.1 | 7 | 9.0 | 26.0 | | 19 | 5.0 | 27.2 | | 21 | 4.3 | 51.173 | 1305.651 | 25.587 | 29.674 | 0.862 | | 0.429 |
| White Blood Cell Counts (×103/ ul) | | 6.0 | 7 | 1.4 | 6.5 | | 19 | 3.1 | 8.0 | | 21 | 3.6 | 29.430 | 433.771 | 14.715 | 9.858 | 1.493 | | 0.236 |
| Red Blood Cell Counts (milli/ul) | | 4.6 | 7 | 0.5 | 4.1 | | 19 | 0.7 | 4.5 | | 21 | 0.5 | 1.879 | 13.901 | 0.940 | 0.316 | 2.974 | | 0.061 |
| Hemoglobin (g/dl) | | 13.1 | 7 | 0.9 | 12.1 | | 19 | 1.7 | 12.6 | | 21 | 1.2 | 6.189 | 83.966 | 3.094 | 1.908 | 1.621 | | 0.209 |
| Mean Corpuscular Volume (fl) | | 84.4 | 7 | 3.5 | 85.2 | | 19 | 4.4 | 83.0 | | 21 | 6.3 | 48.705 | 1199.972 | 24.352 | 27.272 | 0.893 | | 0.417 |
| Mean Corpuscular Hemoglobin (Pg) | | 28.6 | 7 | 1.6 | 29.5 | | 19 | 1.7 | 27.9 | | 21 | 2.5 | 24.427 | 191.035 | 12.213 | 4.342 | 2.813 | | 0.071 |
| Fasting Blood Glucose (mg/dl) | | 109.4 | 7 | 44.4 | 115.4 | | 19 | 32.2 | 107.3 | | 21 | 32.1 | 670.647 | 51231.013 | 335.324 | 1164.341 | 0.288 | | 0.751 |
| HemoglobinA1c % | | 5.4 | 7 | 1.6 | 5.7 | | 19 | 1.6 | 4.8 | | 21 | 0.8 | 9.087 | 71.749 | 4.543 | 1.631 | 2.786 | | 0.073 |
| Serum Cholesterol (mg/dl) | | 165.7 | 7 | 28.4 | 179.9 | | 19 | 36.5 | 180.1 | | 21 | 35.0 | 1216.206 | 53351.028 | 608.103 | 1212.523 | 0.502 | | 0.609 |
| Serum Triglyceride (mg/dl) | | 112.1 | 7 | 76.9 | 98.8 | | 19 | 32.3 | 94.6 | | 21 | 34.9 | 1612.692 | 78632.967 | 806.346 | 1787.113 | 0.451 | | 0.640 |
| High Density lipoprotein (mg/dl) | | 45.0 | 7 | 10.1 | 51.3 | | 19 | 15.4 | 50.9 | | 21 | 9.5 | 221.145 | 6677.494 | 110.572 | 151.761 | 0.729 | | 0.488 |
| Low Density Lipoprotein (mg/dl) | | 85.9 | 7 | 28.3 | 98.4 | | 19 | 33.0 | 99.4 | | 21 | 23.9 | 1031.368 | 35762.632 | 515.684 | 812.787 | 0.634 | | 0.535 |
